# A PD-1-ST2 axis controls Th2 effector function in tissue via a metabolic checkpoint

**DOI:** 10.1101/2023.08.13.553117

**Authors:** Graham A. Heieis, Bart Everts, Craig W. Roberts, Rick M. Maizels, Georgia Perona-Wright

## Abstract

Type 2 immune responses characterise both helminth infections and atopic disease such as allergy or asthma, but a better understanding of the mechanisms that regulate these responses is key to improving therapeutic and vaccination strategies. Immuno-metabolic studies over the last two decades have suggested T cell activation broadly requires rapid increases in glycolysis and oxidative phosphorylation. In contrast, we show that CD4^+^ T helper 2 (Th2) cells activated *in vivo*, using models of helminth infection, do not acquire a glycolytic metabolism. Instead, we show that Th2 cells solely increase their oxidative metabolism, associated with increased fatty acid uptake. Rather than contributing directly to effector function, our data reveal that Th2 cells switch to fatty acid oxidation downstream of PD-1 signalling to promote expression of the IL-33 receptor (ST2). These data provide insight into the spatial regulation of T cell metabolism, and suggest that PD-1 blockade therapies may be effective in Th2 disorders.

## Introduction

T helper (Th)2 cells are required for resistance to helminth infection, affecting nearly 2 billion people, and are also key drivers of atopic disease which is a growing epidemic in western regions^1,2^. As targeting immunometabolism is a promising strategy to treat auto-immunity, cancer and chronic infection, similar principals may be applicable to Th2-associated disease. Furthermore, the signals needed for *in vivo* Th2 differentiation still represent a major gap in our knowledge of CD4^+^ T cell biology. Therefore, understanding the metabolic characteristics and requirements of Th2 cells may yield valuable insights into how Th2 cells develop during infection or disease.

An intimate link exists between metabolic activity in T cells and their effector function. T cell activation triggers a switch from catabolic to anabolic metabolism in order to synthesize necessary macromolecules for DNA synthesis, membrane formation and translation of effector molecules^3^. The importance of metabolic regulation in T cells is highlighted by the association between dysregulated T cell metabolism in disease^4^. Hyper-metabolism in T cells has been shown in patients with or models of multiple sclerosis (EAE)^5,6^, lupus^7^, graft-versus host disease^8^, and obesity^9^. Anergic or exhausted tumour-infiltrating T cells, or those present in viral chronic infection, adopt a hypo-metabolic phenotype, which when reversed is functionally restorative^10,11^.

The majority of metabolic studies in T cells have been done using CD8^+^ T cells or CD4^+^ Th1, Th17 and Treg subsets. In Th1 and Th17 effector cells glycolysis, glutaminolysis and fatty acid synthesis (FAS) are required for differentiation and effector function^6,9,12–14^. Many studies have elucidated that these pathways act through different mechanisms to respectively promote Th1 or Th17 function^5,9,15^. Metabolic activity similarly impacts the Treg-Th17 axis, as blocking glycolysis during Th17 promotes divergent induction of Tregs, which primarily rely on OxPhos fuelled by increased fatty acid oxidation (FAO)^6,16^. Hence, the quality and type of immune response can be altered through manipulating the activity of intrinsic metabolic pathways in CD4^+^ T cells.

In comparison, little work has been done to understand metabolism in Th2 cells. Studies thus far have suggested that glycolysis is tightly paired to Th2 cell activation and function^16–18^. Naïve CD4^+^ T cells polarized *in vitro* under Th2 conditions present with the highest glycolytic rate compared to cells cultured in Th1, Th17 and T_reg_ conditions^6,16^. Pharmacological inhibition of glycolysis with glucose analogue 2-deoxy-*d*-glucose (2DG) completely abrogates IL-4 production indicating the pathway is needed for optimal Th2 function^18^. Impairing the ability of Th2 cells to engage glycolysis *in vivo* using mice with a T cell specific knockout of the environmental sensor mammalian target of rapamycin (mTOR), a central regulator of T cell metabolism, also translated to reduced cytokines, tissue-infiltrating cells, and pathology during allergic challenge^18^. Combined these data have been interpreted to mean that a high glycolytic rate is an inherent feature of Th2 cells, but this interpretation has been derived mainly from *in vitro* models. The metabolic properties of Th2 cells have yet to be sufficiently characterized directly from an *in vivo* source during an ongoing immune response.

In this study, we therefore aimed to determine a metabolic phenotype for Th2 cells generated *in vivo* and its relevance to Th2 effector function. We describe Th2 cells as being a poorly glycolytic subset that maintain elevated OxPhos due to a preferential use of fatty acids. We further propose metabolism in Th2 cells is dictated by Programmed Death Protein-1 (PD-1) signalling and functions to promote ST2 induction for the detection of the tissue alarmin IL-33.

## Results

### Distinct Th2 metabolism *in vivo* compared to Th1 and *in vitro* polarised Th2 cells

Using IFNγ or IL-4 transcriptional reporter mice, we first assessed metabolism of sorted cytokine^+^ Th1 or Th2 cells polarized from naïve splenocytes *in vitro* (Figure 1A). In agreement with previous findings, Th2 cells trended towards higher glycolytic gene expression (Figure 1B) and had a significantly higher extracellular acidification rate (ECAR), representative of glycolysis (Figure 1C). However, the oxygen consumption rate (OCR) was similarly increased in Th2 cells so that the OCR/ECAR ratio was comparable to Th1 counterparts, suggesting a parallel increase in mitochondrial OxPhos (Figure 1C). Oligomycin injection, used to induce maximal glycolysis, additionally had little effect on the ECAR of either subset (Figure 1C). Therefore, despite elevated metabolic activity in Th2 cells compared to Th1, T_eff_ cells activated *in vitro* are highly and maximally glycolytic.

**Figure 1:**
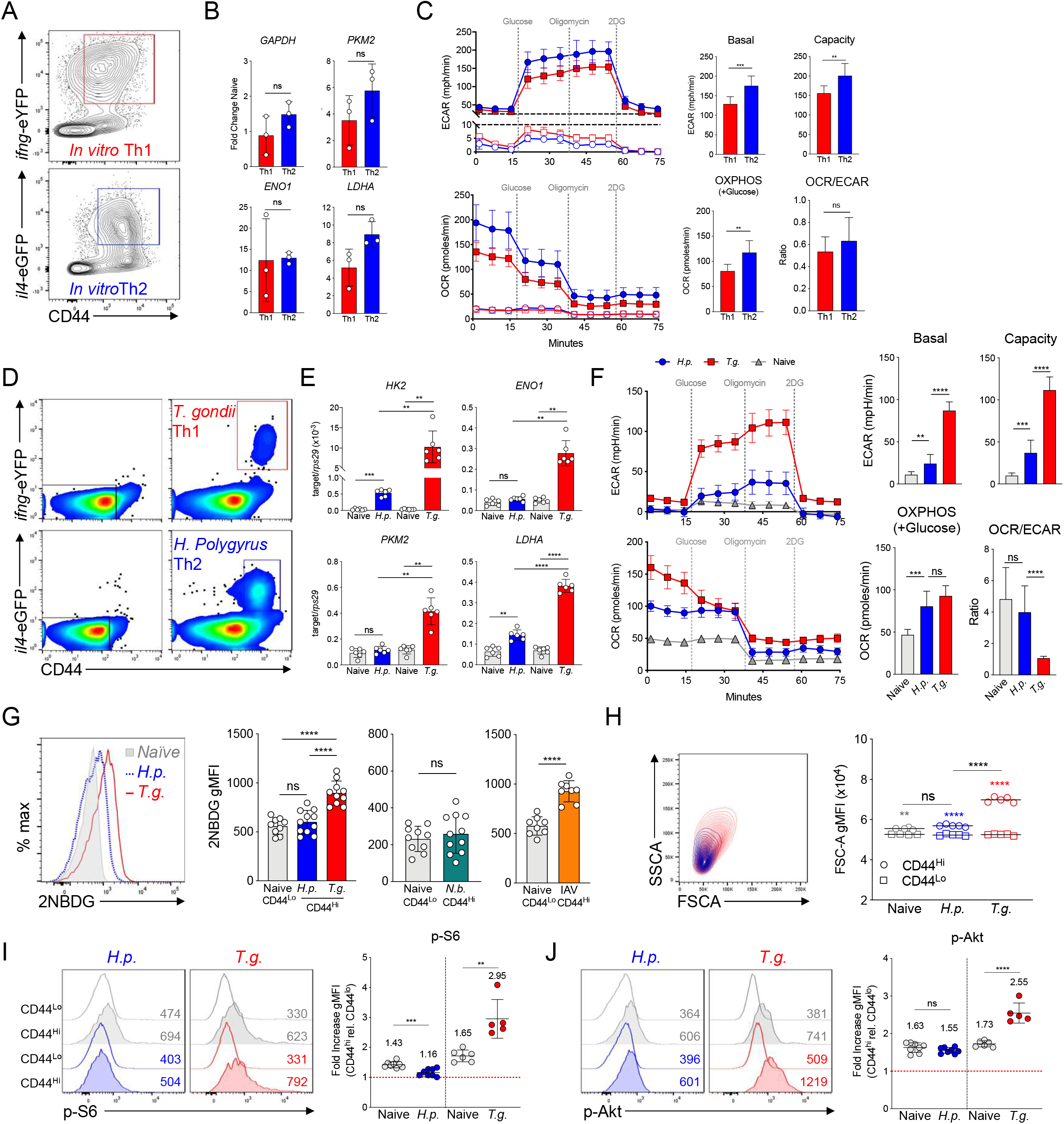
Th2 cell metabolism is distinct ex vivo from Th1 or in vitro-polarized CD4^+^ T cells. (A-C) Metabolic characterization of Th1 and Th2 cells activated *in vitro.* (A) Representative reporter expression and sort gates for Th1 or Th2 polarized CD4^+^ splenocytes, from Great (*ifng-* eYFP) or B6.4get (*il4-*eGFP) mice respectively, 4 days after activation. (B) Glycolytic enzyme gene expression in cytokine^+^ cells: Hexokinase 2 (*Hk2*), glyceraldehyde 3-phosphate dehydrogenase (*Gapdh*), enolase 1 (*Eno1*), Pyruvate kinase M2 (*Pkm2*); data points represent independent experiments with mean and SD. (C) Real-time glycolytic (extracellular acidification) rate (ECAR) and oxygen consumption rate (OCR) of sorted Th1 and Th2 cells using the Seahorse glycolytic stress test, pooled from 3 independent experiments. (E-F) Reporter mice were infected with either *T. gondii* for 7 days or *H. polygyrus* for 14 days. (E) Representative reporter expression and sort gates for *in vivo* activated Th1 or Th2 cells, or naïve CD4 T cells from uninfected controls, isolated from the mesenteric lymph. (E) Glycolytic enzyme gene expression from MLN sorted cytokine^+^ cells; data points represent individual mice, pooled from 2 experiments of 3 mice each with mean and SD shown. (F) Extracellular flux analysis of *ex vivo* sorted cells using the Seahorse glycolysis stress-test; 3 experiments combined with cells pooled from 7-10 mice per experiment and plated as technical replicates. (G) C57BL/6 mice were injected with 2NBDG during infection and CD4^+^ T cells from the draining LN were measured for uptake. Cells were taken either from the MLN from Tg or Hp infected mice, or the mediastinal LN of mice infected 7 days with influenza A virus or 9 days with *N. brasiliensis*; mean and SD of individual mice plotted from pooled data of 2-3 experiments with 2-4 mice. (H) Representative forward and side scatter (FSCA/SSCA) profile for CD4^+^CD44^Hi^ cells from the MLN of Tg or Hp infected mice, and gMFI of FSCA comparing CD44^Lo^ and CD44^Hi^ cells from naïve and infected mice. (I,J) Representative histograms with corresponding gMFI for phospho-S6 (I) and phospho-Akt (J) detection and normalized ratio of the CD44^Hi^ compared to CD44^Lo^ population; data pooled from 2 experiments with 2-4 mice each. Statistics analysed by student’s two-tailed t-test or one-way ANOVA: ****p<0.0001, ***p<0.001, **p<0.01, *p<0.05, ns = not significant.

To question whether Th2 cells also had a superior glycolytic rate *in vivo*, we infected reporter mice with *Toxoplasma gondii* (Tg) or *Heligmosomoides polygyrus* (Hp), two enteric parasites that respectively induce robust Th1 or Th2 responses in the draining mesenteric lymph node (MLN) (Figure 1D). Sorted *ifng*^+^ Th1 cells displayed significantly increased glycolytic gene expression compared to strain matched naïve CD4^+^CD44^Lo^ cells from uninfected mice (Figure 1E). In contrast, sorted *il4*^+^ Th2 cells showed a significantly smaller increase in *Hk2* and *Ldha* expression, while *Eno1* and *Pkm2* remained comparable to naïve cells, suggesting Th2 cells have a reduced glycolytic capacity compared to Th1 cells *in vivo* (Figure 1E). Extracellular flux analysis of sorted cells from the MLN indeed revealed a distinctly lower glycolytic rate in Th2 compared to Th1 (Figure 1F). Th2 glycolysis actually appeared to more closely resemble that of naïve cells, although a small significant increase was observed.

Th1 cells also initially displayed a high OCR in comparison to Th2 cells from infected mice when glucose was absent (Figure 1F). However, the provision of glucose led to a similar OCR in both subsets due to a drop by Th1 cells (Figure 1F). Consequently, Th1 cells had a significantly depreciated OCR/ECAR ratio, whereas Th2 cells maintained a ratio matching naïve CD4^+^ cells (Figure 1F). Furthermore, while adding glucose caused Th1 cells to reduce their OCR it had no impact on the OCR of Th2 cells, implying that oxidative metabolism of helminth activated Th2 cells may be less dependent on glucose availability. A similar metabolic phenotype was still observed in sorted Th2 cells after depleting Tregs during infection, or from Hp resistant BALB/c reporter mice, suggesting low glucose metabolism is an intrinsic property of Th2 cells (Figure S1). Altogether, these data demonstrate a discrepancy between the metabolism of *in vitro* and *in vivo* activated Th2 cells. Whereas Th1 and Th2 cells are highly glycolytic following *in vitro* polarization, as are Th1 when activated *in vivo*, Th2 cells responding *in vivo* to parasitic infection remain glycolytically inert and instead selectively increase oxidative metabolism.

### Restricted glucose uptake and mTOR activation in Th2 cells *in vivo*

We next sought to identify at what level the low rate of glycolysis was regulated in Th2 cells during infection. Infected C57BL/6 mice were injected with the fluorescent glucose analogue 2NBDG to assess glucose uptake *in vivo*. Activated T cells from Hp infection acquired glucose at a similar rate to naïve cells from uninfected controls, whereas Th1 cells had a significant increase in 2NBDG uptake (Figure 1G). This observation was replicated by T cells from the lung-draining mediastinal lymph node (MdLN) using influenza A virus (IAV) or *Nippostrongylus brasiliensis* (Nb) as alternative Th1 or Th2 driving infections, respectively (Figure 1G).

We noted that activated T cells from Hp infection had markedly reduced cell, measured by forward-scatter, compared to those from Tg infection (Figure 1H), and hypothesized these findings would be consistent with reduced mTOR activation^18,19^. In CD44^Hi^ T cells from Tg infection, elevated mTOR signalling was evident by increased phosphorylation levels S6 and Akt, respective mTORC1 and mTORC2 targets, compared to CD44^Hi^ cells from unchallenged animals (Figure 1I,J). Conversely Hp infection did not lead to an increase in p-Akt beyond that seen in CD44^Hi^ cells from naïve controls, and resulted in a significant decrease in p-S6 (Figure 1I,J). Thus, reduced glucose uptake, and consequently the low glycolytic phenotype of Th2 cells, could be due to a lack of downstream mTOR signalling.

### Th2 metabolism is dissociated from cytokine production in the tissue

It has recently been appreciated that cytokine production by Th2 cells is temporally regulated between lymph node priming and arrival at the peripheral infection site, in part through exposure to alarmins such as IL-33^20,21^. While investigating potential factors that could drive metabolic alterations in Th2 cells, we found that IL-33 receptor (IL-33R/ST2) expressing cells from the MLN of Hp infected mice selectively increased uptake of the fluorescent fatty acid BODIPY FL C16 (palmitate) (Figures 2A). Interestingly, this increase was not dependent on IL-33, as IL-33 treatment *ex vivo* increased cytokine production but not FA uptake (Figures 2A & S2A).

**Figure 2:**
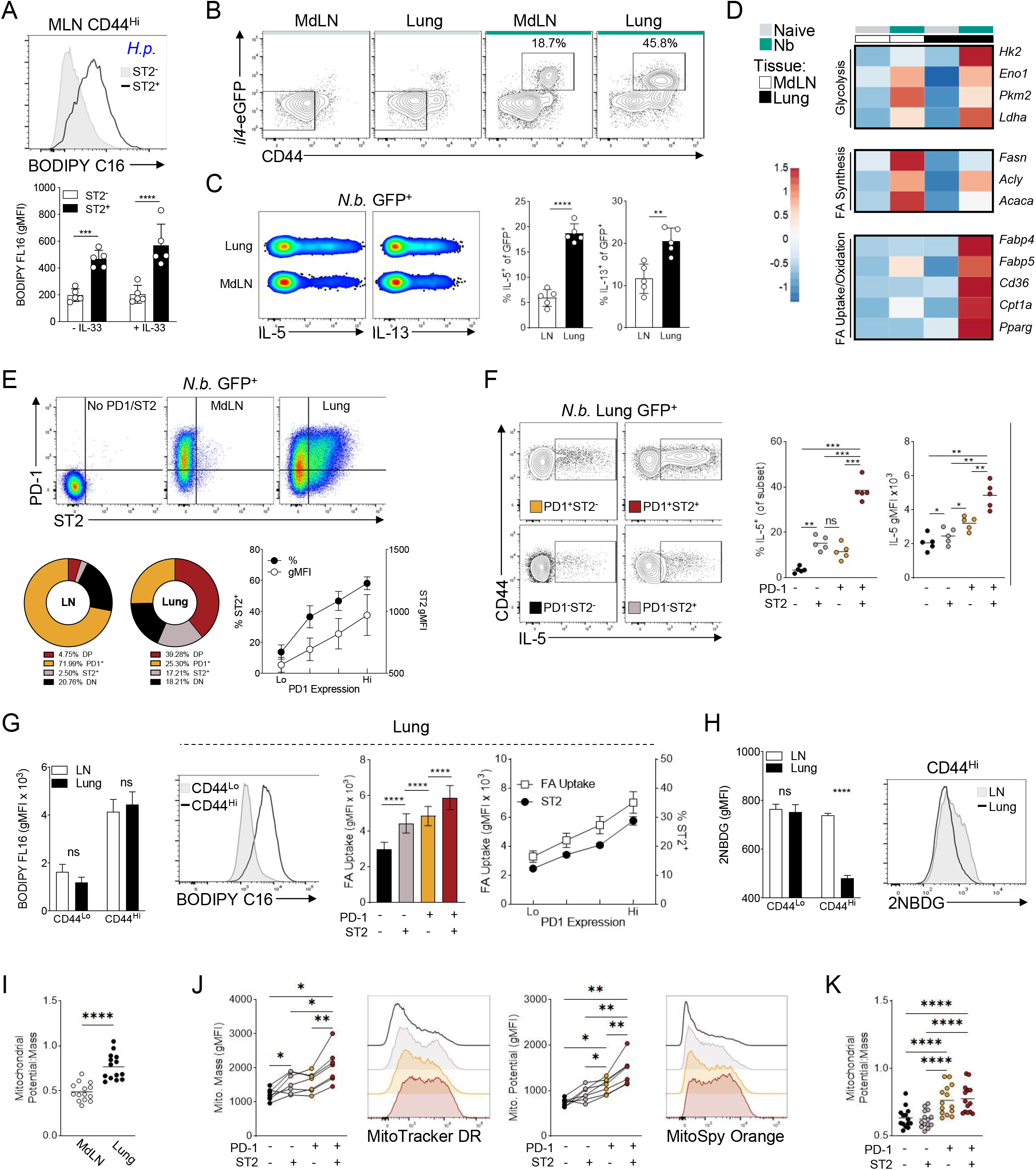
PD-1^+^ST2^+^ Th2 cells in the tissue have enhanced effector function and mitochondrial fitness associated with lipid uptake. (A) Representative histogram and quantification of BODIPY FL C16 uptake in sorted CD4^+^CD44^Hi^ T cells isolated from the MLN of Hp infected mice and stimulated overnight with or without recombinant IL-33; data points are technical replicates of cells pooled from 6-10 mice, representative of 3 experiments. (B) Representative reporter expression and example gating for TCR+CD4+ T cells from the MdLN of Nb infected mice. (C) Intracellular cytokine staining comparing expression of IL-5 and IL-13 between *il4-*eGFP^+^ T cells in MdLN or lung; data are individual mice showing mean and SD, representative of 3 experiments. (D) Heatmap displaying relative gene expression to housekeeping gene of metabolic targets in sorted Th2 or naïve CD44Lo cells as shown in (B), created using ClustVis webtool; data represent the mean of 3 mice per group, with similar results obtained from 2 independent experiments. (E) Co-expression of ST2 with PD-1 in the lung compared MdLN, distribution showing the mean population frequency, and mean ST2 expression associated with PD-1 expression quartiles; pooled from 3 experiments with 6-7 mice. (F) Intracellular IL-5 staining according to PD-1 and ST2 expression; representative of 2 experiments with 5-7 mice. (G, H) Metabolite uptake in CD4^+^ populations isolated by negative selection from the MdLN or lung of Nb infected C57BL/6 mice for BODIPY FL C16 (G) and 2NBDG (H); mean and SD shown for technical replicates from a pool for 4-6 mice, representative of 3 experiments. (I) Ratio of MitoSpy Orange (mitochondrial potential) to MitoTracker DeepRed (mass) staining in GFP^+^ T cells from the MdLN or lung of B6.4get mice infected with Nb. (J) MitoTracker Deepred and MitoSpy Orange staining according to PD-1 or ST2 expression in lung Th2 cells, and the ratio of mitochondrial potential to mass (K). Statistics performed using students’ two-tailed t-test (A, C, G-I), one-way ANOVA (F) or RM one-way ANOVA when comparing populations within a matched sample (G, J, K): ****p<0.0001, ***p<0.001, **p<0.01, *p<0.05, ns = not significant.

To further analyse Th2 metabolism in the context of tissue-based signals, we used Nb infection to obtain *il4-*eGFP^+^ cells from the MdLN and lung, of which the latter is a site for terminal Th2 differentiation^20^. Accordingly, the lung harboured a greater frequency of GFP^+^ cells, which secreted more IL-5 and IL-13 compared to those from the MdLN (Figure 2B,C). Expression of glycolytic genes in total GFP^+^ cells was increased compared to naïve CD44^Lo^ cells, but similar between sites during infection (Figure 2B,D). Conversely, genes for FAS were significantly increased in Th2 from the MdLN compared to both naïve CD44^Lo^ or Lung GFP^+^ Th2 cells (Figure 2D). Lung Th2 cells, however, uniquely increased expression of genes involved with FA uptake, as well as *Pparg*, a transcriptional regulator of lipid metabolism (Figure 2D). Similar expression patterns were also observed using a model of house-dust mite allergy (Figure S2C). We tested the requirement for environmental glucose or lipids for Th2 cytokine production by directly stimulating lung homogenate from infected mice. Despite the low glucose metabolism exhibited by Th2 cells, IL-4, IL-5 and IL-13 expression all demonstrated an acute sensitivity to glucose availability, as well as to the hexokinase inhibitor 2DG (Figure S2C), which may be a result of blocked translation rather than specific impairment of Th2 function^22^. In contrast, cytokine expression remained comparable to control conditions in the presence or absence of FA, or inhibitors of lipolysis or FAO (Figure S2D). These findings therefore illustrate Th2 cells undergo location-dependent changes in metabolism, though these metabolic changes do not necessarily provide a direct contribution to cytokine expression.

### ST2^+^ Th2 enriched in the tissue exhibit metabolic properties in line with PD-1 signalling

We considered the possibility that FA metabolism had an indirect role by supporting terminal Th2 differentiation in the tissue. Accordingly, analysis of ST2 on *il4*-eGFP^+^ cells during Nb infection revealed its robust expression on lung Th2 cells, but its near-absence on those remaining in the MdLN (Figure 2E). PD-1 Ligand 1 (PD-L1) can promote Th2 responses in the lung of mice infected with Nb, but how it signals to support Th2 function is not yet known^23^. Interestingly, a metabolic switch from glycolysis to lipolysis is a hallmark of PD-1 associated T cell exhaustion^24^. Combining these concepts with our data that ST2^+^ cells have a selectively altered metabolism, we investigated PD-1 signalling as a possible regulator of metabolism in ST2^+^ Th2 cells.

Unlike ST2, surface PD-1 was abundant on cells from both sites (Figure 2E). However, ST2 had stringent co-expression with PD-1, with up to 80% of ST2^+^ cells also expressing PD-1 (Figure 2E). PD1^+^ST2^+^ Th2 cells were the most potent produces of IL-5 and IL-13, further supporting these cells as more terminally differentiated effectors (Figure 2F & S2E). Total CD44^Hi^ T cells from Nb infection had significantly increased FA uptake compared to naïve cells from control mice, irrespective of being isolated from the MdLN or Lung, but within the lung PD1^+^ST2^+^ cells demonstrated the greatest uptake (Figure 2G). In contrast to FA uptake, CD44^Hi^ cells from the lung showed significantly reduced 2NBDG uptake compared to those from the MdLN, indicating a further shutdown in glucose metabolism upon tissue-entry (Figure 2H). In support of FA being shuttled into OxPhos, lung Th2 possessed a higher ratio of mitochondrial potential to mass compared to the MdLN (Figure 2I). Amongst Th2 from the lung, PD1^+^ST2^+^ further had the greatest mitochondrial mass and potential, but also the greatest potential to mass ratio. Thus, Th2 cells, from the lung in particular, possess a metabolic profile consistent with that described downstream of PD-1 signalling.

To more deeply interrogate metabolic differences, we used a metabolic spectral flow-cytometry panel to compare enzyme/transporter expression between T cell populations from the MdLN or lung^25^. MdLN and lung samples from individual mice were CD45-barcoded and pooled prior to staining to eliminate differences driven by cell composition (Figure S3A). Comparing tissues from infected mice, T cells from the lung demonstrated a marked decrease in all measured targets regardless of activation state, such that MdLN and lung cells clustered completely separately (Figure 3A, B & S3B). Manual gating confirmed that both CXCR5^+^ T follicular helper cells (Tfh) and GATA3^+^ Th2 cells in the MdLN had globally increased metabolic protein expression compared to naïve cells (Figure 3C). Th2 cells also showed overall higher expression than Tfh, with the notable exception of glucose transporter GLUT1. A specific increase in Tfh glucose metabolism was supported by higher 2NBDG, indicating glucose metabolism may be a bifurcation between Tfh and Th2 (Figure 3D).

**Figure 3:**
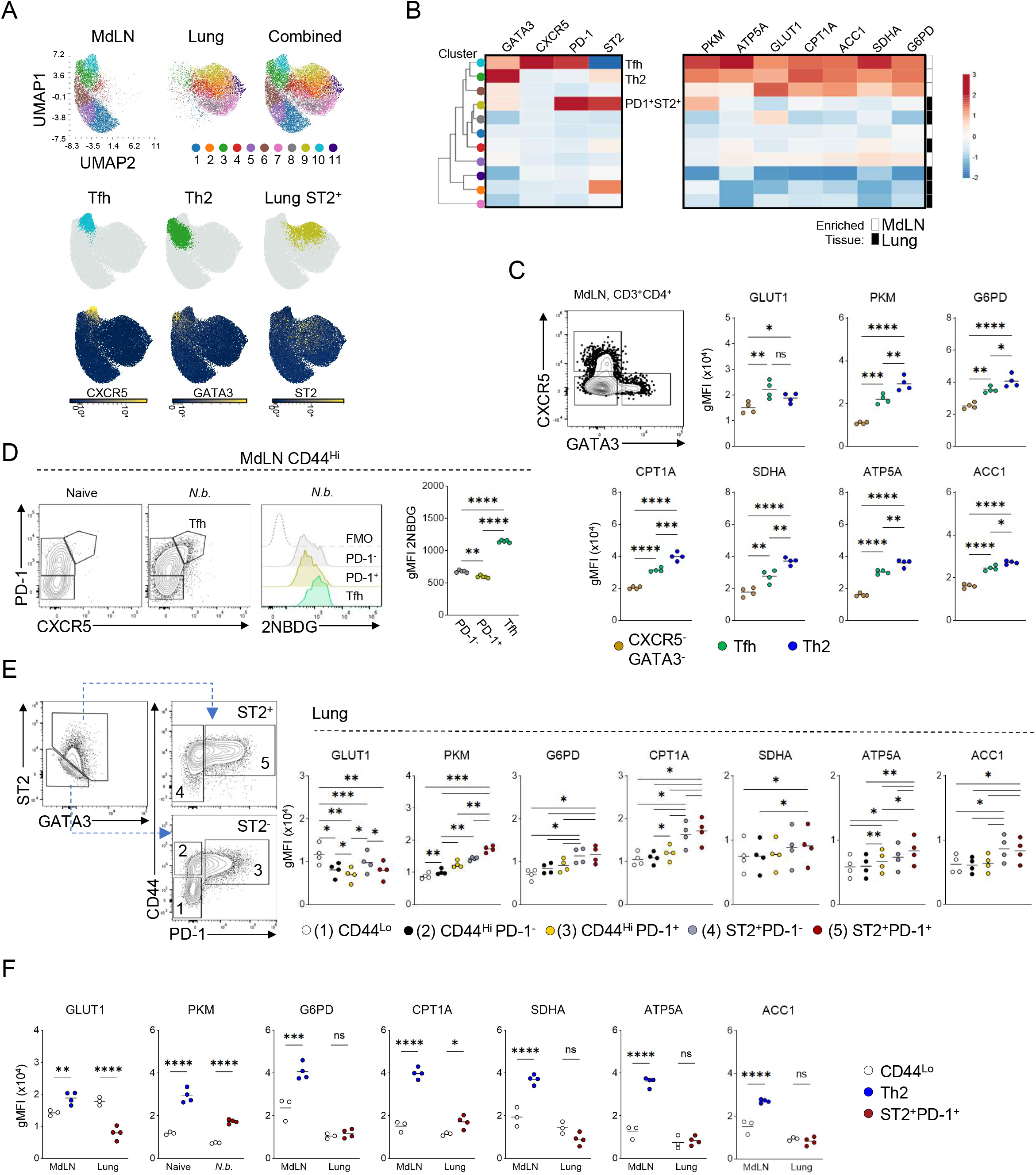
Metabolic flow cytometry analysis of lymph node and lung Th2 cells. (A) UMAP and phenograph clustering using metabolic and phenotypic markers of total CD3+CD4+ T cells from the MdLN or Lung of Nb infected C57BL/6 mice performed using OMIQ using subsampled data from 4 mice. (B) Heatmap showing median expression of phenotypic and metabolic markers for corresponding clusters in (A), created using ClustVis. (C) Manual Tfh and Th2 gating and associated metabolic target expression for the MdLN from mice used in (A). (D) *ex vivo* 2NBDG uptake of CD44^Hi^ T cell populations from the MdLN of infected mice; representative of 3 experiments using cells pooled from 4-6 mice. (E) Manual gating and analysis for metabolic protein expression of lung CD4^+^ T cell populations during Nb infection for mice used in (A). (F) Comparison between GATA3^+^ Th2 from the MdLN or PD-1^+^ST2^+^ cells from the lung of infected mice with tissue matched naïve CD44^Lo^ T cells; representative of 2 experiments. Statistics calculated using RM one-way ANOVA (C-E) or two-way ANOVA comparing means within tissue (F): ****p<0.0001, ***p<0.001, **p<0.01, *p<0.05, ns = not significant.

Analysis of the lung revealed a limited increase in metabolic markers in ST2^+^ cells over other populations, apart from GLUT1, which supports the notion of reduced glucose metabolism in the lung as seen by 2NBDG uptake (Figure 3E & 2H). When comparing across tissues, CD44^Lo^ cells from uninfected mice appeared relatively comparable indicating changes in metabolic expression are not solely driven by changes in environment (Figure 3F). Importantly, in lung PD1^+^ST2^+^ cells, only CPT1A and PKM were increased relative to naïve CD44^Lo^ cells, whereas GLUT1 was again the only significantly reduced marker (Figure 3F). These observations, altogether, suggest that lung ST2^+^ Th2 cells maintain the ability to perform increased FAO while shutting down glucose metabolism.

### PD-1 signalling induces ST2 expression in a metabolically dependent manner

We hypothesized PD-1 initiates a metabolic switch permissive for ST2 induction in peripheral tissues. In support of this hypothesis, *ex vivo* stimulation of sorted *il4-*eGFP^+^ in the presence of αCD3 and PD-L1-Fc led to a stark increase in the frequency and intensity of ST2 expression (Figure 4A). Increased ST2 was not observed with either αCD3 or PD-L1-Fc stimulation alone, indicating both T cell stimulus and PD-1 ligation are required for optimal ST2 induction. Interestingly, dual stimulation also significantly reduced the blasting profile and *il4* expression in responding cells, despite increasing ST2 (Figure 4B,C). Paralleling these data, we also observed decreased size, GATA3 and Ki67 in activated cells from the lung compared to the MdLN during Nb infection (Figure 4D-F). Lung Th2 cells, however, possessed increased granularity (measured by side-scatter), a feature commonly associated with high cytokine producing cells (Figure 4D).

**Figure 4:**
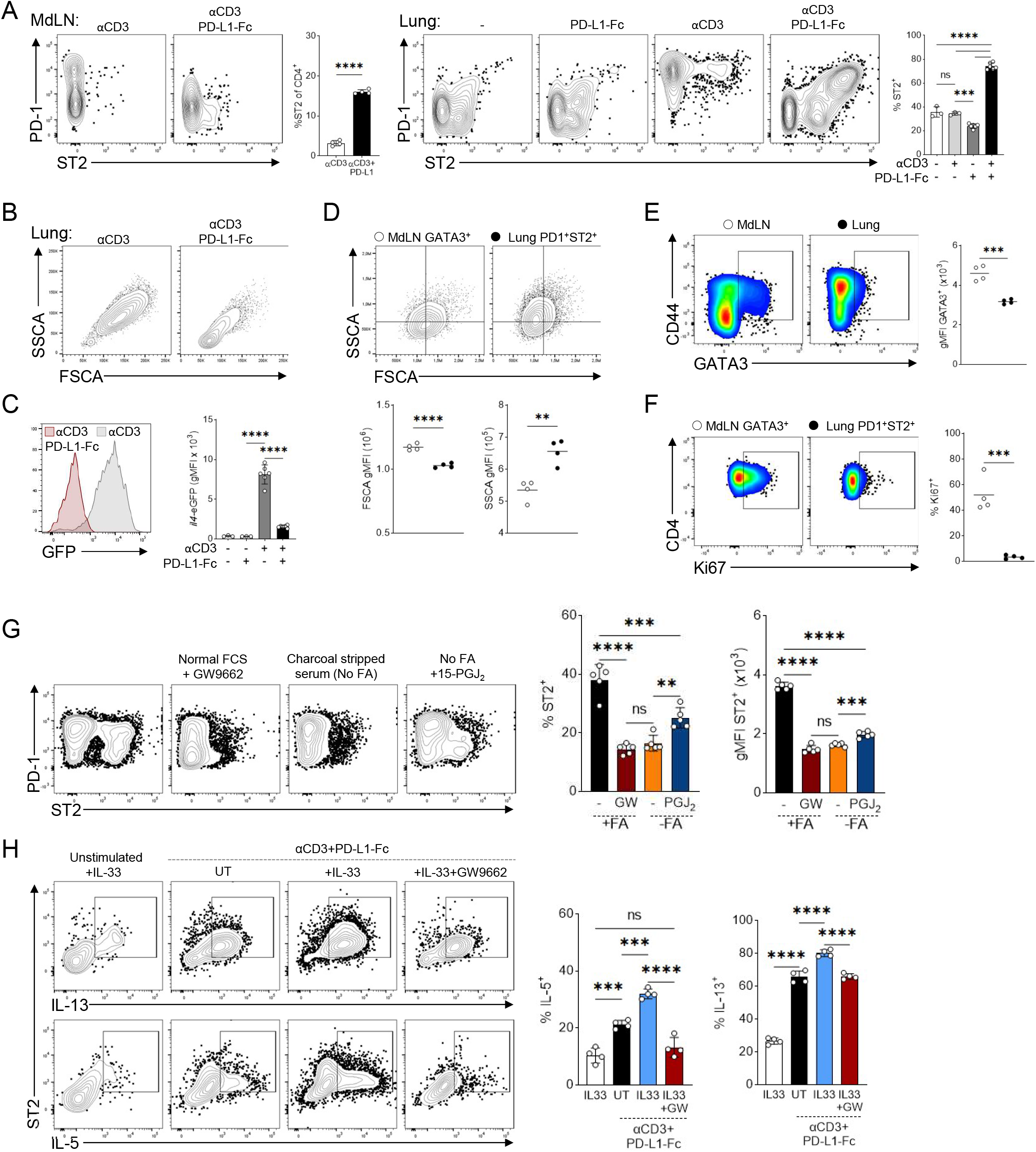
PD-1 signalling promotes ST2 expression in a metabolically dependent manner. (A) PD-1 and ST2 expression of sorted *il4-*eGFP^+^ cells from Nb infected mice following 3 days of culture with either CD3 or PD-1 stimulation; representative of 5 experiments, showing technical replicates from a pool of 7-10 mice. (B) Example plots showing blasting profile of ex vivo stimulated Th2 cells with or without PD-L1-Fc treatment, sorted form the lung of infected mice. (C) *il4*-eGFP expression of sorted and *ex vivo* stimulated Th2 cells after CD3 or PD-1 stimulation. (D) Blasting profile of indicated CD4+ populations assessed directly ex vivo from the MdLN or Lung during Nb infection. (E) Comparative GATA3 expression between total CD4^+^ T cells from the infected MdLN or lung. (F) Corresponding Ki67 staining of indicated CD4^+^ T cell populations; Representative of 3 experiments (E-F), showing mice as individual data points. (G) Sorted Th2 cells cultured as in (A) in the presence or absence of PPARγ inhibitor GW9662 in media with normal FCS or PPARγ activator 15-PGJ_2_ in media with charcoal-stripped serum depleted of fatty-acids and lipids; comparable results achieved from 2 independent experiments, showing technical replicates from a pool of 7-9 mice. (H) Sorted cells stimulated as in (A) with or without recombinant IL-33 or PPARγ inhibition and assessed for intracellular cytokines after PMA/Ionomycin stimulation; data from one experiment with technical data points from 8 mice pooled. Statistics performed using students’ two-tailed t-test when comparing 2 groups or one-way ANOVA for comparisons between more than 2 groups: ****p<0.0001, ***p<0.001, **p<0.01, *p<0.05, ns = not significant.

Previous studies have demonstrated ST2 expression is regulated by the transcription factor PPARγ, a driver of FAO^26,27^. Accordingly, PPARγ inhibition with GW9662 severely impaired ST2 induction driven by PD-1 (Figure 4G). To determine if ST2 expression downstream of PD-1 is dependent on FA, we performed *ex vivo* stimulations using media supplemented with charcoal-stripped serum that lacks FA. Remarkably, without FA present, the increase in ST2 driven by PD-1 signalling was completely blocked (Figure 4G). To address if FA-free conditions impaired ST2 as a consequence of failed PPARγ activation, we added the PPARγ agonist 15-PGJ_2_ to Th2 cells re-stimulated in FA-free media. This achieved partial restoration, indicating that PPARγ can partially promote ST2 expression independent of FA metabolism, yet maximal expression can only be achieved with FA present (Figure 4G). Thus, altogether our data align with a model in which tissue-localized PD-1 signalling permits IL-33 responsiveness in a metabolically-dependent manner, while limiting untargeted cytokine production from immigrating, LN-primed Th2 cells (Figure 5).

**Figure 5:**
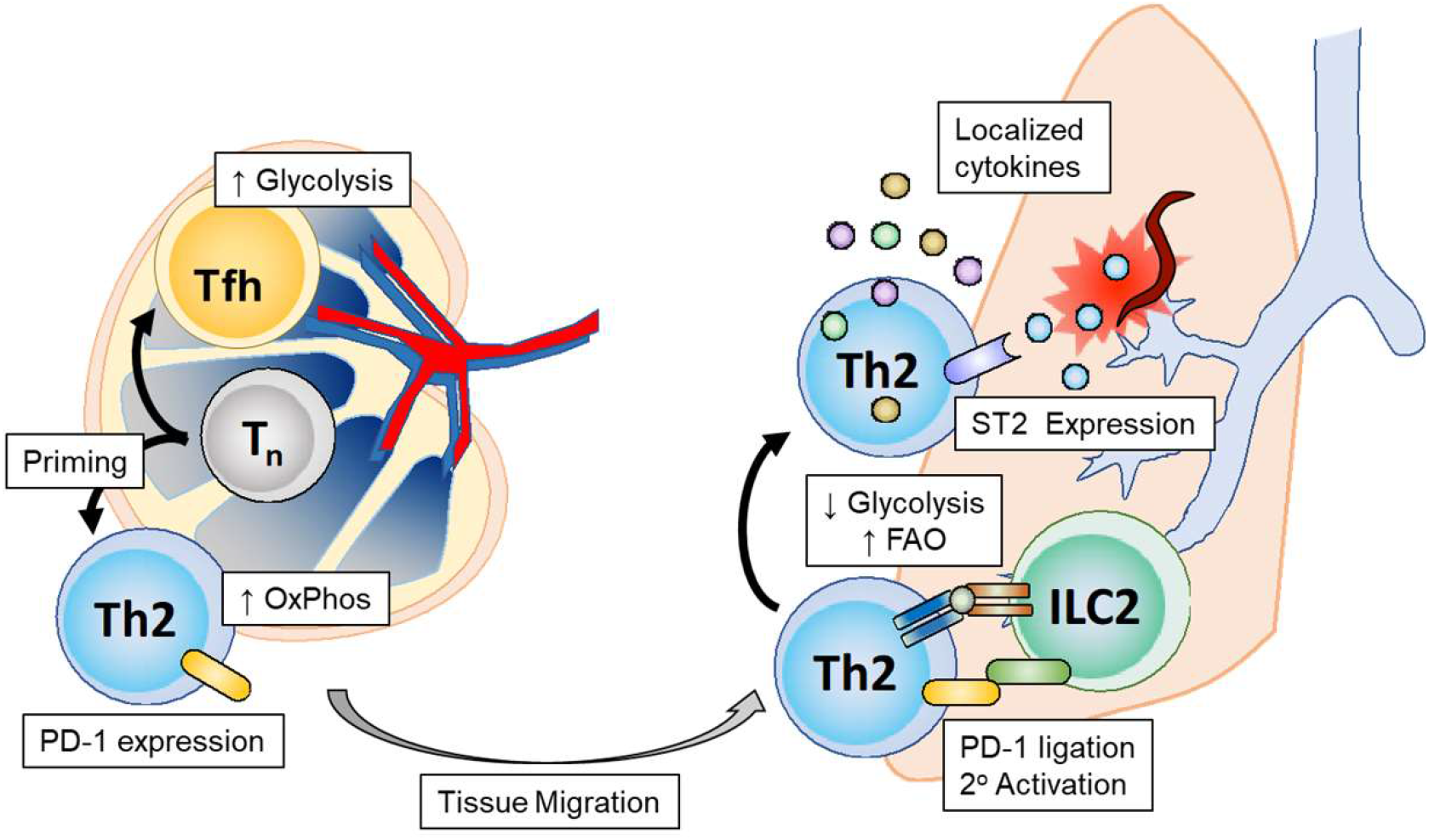
Proposed model for PD-1 regulation of tissue Th2 metabolism and immunity. During initial T cell priming, Th2 cells elevate mitochondrial metabolism downstream of TCR signalling, but independent of glucose metabolism. In contrast glycolysis represents a bifurcation between Th2 and Tfh, whereby Tfh retained in the lymph node become highly glycolytic. Primed Th2 acquired PD-1 expression before emigration from the lymph node and encounter PD-L1 signals upon entry into the tissue site, possibly from resident ILC2 populations. PD-1 signalling triggers a further shutdown in glucose metabolism while maintaining capacity of FAO and mitochondrial metabolism, thus restraining untargeted cytokine production. This metabolic switch simultaneously promotes expression of the IL-33 receptor ST2 such that when damage-associated signals are present, cytokine expression is re-initiated at the appropriate time and location. Thus, we ultimately propose PD-1 as a tissue-based metabolic checkpoint to control unwanted Th2 responses while permitting timely cytokine production when required.

## Discussion

It is widely acknowledged that Th2 cells are a highly glycolytic subset, requiring glucose metabolism for differentiation and cytokine production^4,28–30^. We and others have shown that this is true *in vitro*^6,16,18^. However, there is a broad gap in our understanding of how metabolism is regulated *in vivo*. By analysing cytokine-expressing T cells directly from the draining lymph nodes of infected mice, we find that Th2 cells are poorly glycolytic. Instead, our data suggest that Th2 cells rely heavily on glucose-independent mitochondrial metabolism, with fatty acids as a dominant fuel source.

Our findings are supported by other reports alluding to preferential FAO in Th2 cells^31^. Sequencing studies have correlated FAO genes to pathogenic Th2 cells in eosinophilic esophagitis^32^ and allergic asthma^33^. Additionally, much recent work has been done on the PPAR family of transcription factors, master regulators of lipid metabolism. PPARγ in particular has been consistently linked to Th2 differentiation^26,31,34^. In macrophages and ILC2 PPARγ has been shown to directly promote FA uptake and oxidation^27,35,36^, and could play a similar role in Th2 cells. While less is known about the other PPARs in T cells, PPARα, a driver of FAO in adipocytes^37^, also appears to be uniquely expressed and required in Th2 cells^38^. Interestingly, FAO appears to be a common trait amongst immune populations in type 2 immunity, including alternatively activated macrophages, ILC2s and likely Th2-priming DCs^34,36,39^.

In this study, we focused on the metabolic requirements of Th2 cells for effector function, rather than differentiation. By preventing the use of fatty acids or lipids during *ex vivo* stimulation, we found cytokine production was acutely independent of fatty acid metabolism. Although glucose was found to be required, it has been shown that T cells have a high translational dependence of glucose-derived ATP, which could partly explain the discrepancy with low glucose metabolism in Th2 cells^22^. Interestingly, however, Tfh cells in the draining LN displayed significantly increased glucose uptake and GLUT1 expression. Increased glycolysis may then be required for T cells functioning within the follicular zone, perhaps due to its hypoxic nature^40^. Intriguingly, Th2 glucose metabolism dropped even further in the lung, which may be partially due to the low glucose availability there^41,42^. Generally, however, we found nearly all metabolic targets to be decreased relative to LN Th2 cells which could also be linked to impaired translational capacity from environmental restrictions.

Our data instead led us to the discovery that lipid metabolism in Th2 cells is a requisite for IL-33 receptor expression in peripheral tissues, and that this was controlled by PD-1 signalling. PD-1 has previously been identified as a Th2-promoting factor during Nb infection^23^. Separately, PD-1 ligation has been shown to suppress glycolysis while diverting fatty acids towards mitochondrial oxidation^24^. Together, these discoveries prompted us to investigate the link between PD-1 and Th2 metabolism and we thus elucidated their combined impact on ST2. We and others have also found that ST2 expression was localized mainly to the peripheral tissue as opposed to priming site^20^, aligning with the proposal that ILC2s, abundant in the lung but rare in the LN^43^, are the major source of PD-1 ligands^23^. As PD-1 can also interfere with mTOR signalling, a key promoter of translation in T cells^44,45^, this could further explain the consistent reduction in protein expression in lung Th2 cells compared to those in the LN. Yet, lung Th2 cells still retained elevated lipid uptake and CPT1A expression suggesting a dominant reliance on FAO, further supportive of a PD-1 driven metabolic phenotype^24^.

IL-33 plays a critical role in the pathology of allergic and asthmatic inflammation, and therefore IL-33 targeted therapies are being actively trialled for their clinical benefit^46^. However, little is currently known about what regulates expression of its receptor ST2, which is curiously only expressed *in vitro* after multiple rounds of stimulation and independent of Th2 cytokines^47,48^. PPARγ activation has been connected to ST2 expression in Th2 cells, as well as a subset of GATA3^+^ Tregs in adipose tissue^26,49^. More recently in ILC2, the ST2 promoter region has been identified as a direct binding target of PPARγ^35,50^. In agreement with our proposed model that PPARγ activation at peripheral sites drives ST2 expression, PPARγ deficiency led to a defect in cytokine production in the lung, but not LN of HDM-challenged mice, due in part to a reduction in ST2^+^ cells^26^. Following HDM-challenge a large ST2^+^ population, however, remained amongst PPARγ-deficient T cells, indicating ST2 may also be expressed via PPARγ independent mechanisms. Similarly, PPARγ-null mice exhibit only a partial decrease in ST2^+^ Tregs in visceral adipose tissue^49^. In our hands, PPARγ was also required for ST2 induction following PD-1 ligation, yet activating PPARγ in the absence of lipids was not sufficient for full restoration. We therefore suggest fatty acid metabolism has an additional role in determining ST2 expression. Indeed, a direct association between metabolism and ST2 expression has been defined in ILC2^51^. Glycolytic activity was shown to inhibit ST2 transcription through histone methylation of the gene locus. Whether FAO played a role was not investigated, but ILC2 expansion has been demonstrated to rely on FAO during helminth infection^36^. We also observed a reduction in glucose metabolism in lung Th2 cells compared to the MdLN, indicating a similar metabolic-epigenetic link could potentially contribute to ST2 expression here.

It is intriguing as to why distinct CD4^+^ subsets, that are seemingly all proliferative effector cells producing cytokine, would have distinct metabolic properties. The importance of the nutrient environment in dictating cellular metabolism has become increasingly realised, and therefore results may be biased by the tissue in question^52–54^. However, we still observe a stark contrast between Th1 and Th2 cells glycolysis at the same priming site, supporting and inherent difference between subsets. This follows an increasingly apparent dichotomy between the cellular metabolism of type 1 and type 2 immunity, which may also be influenced by changes in nutrient availability in connection with whole-body metabolism. Viremia or bacteraemia frequently cause hyperglycaemia in mice and patients^55^. Interestingly, the inverse is true of helminth infection^56^. Therefore, metabolic responses in immunity may be linked to the evolution of the host-pathogen arms race. Here we demonstrate one possible outcome of how Th2 responses adapt to a potentially low glucose environment.

## Methods

### Mice

C57BL/6 mice were obtained in Canada from Jackson Laboratories. B6.4get and Great mice were ordered initially from Jackson Laboratories then bred in-house under specific pathogen-free conditions at the University of British Columbia. In the United Kingdom, C57BL/6 mice were obtained from Envigo and B6.4get mice were generously provided by Dr. John Grainger at the University of Manchester and bred in-house at the University of Glasgow in conventional facilities, maintained in IVC cages. All animal work was conducted in accordance with the guidelines set by the University of British Columbia Animal Care Committee and the Canadian Council of Animal Care, or with the approval of the University of Glasgow Animal Welfare and Ethics Review Board and done under licensing issued by the UK Home Office. Mice were infected between 6-12 weeks of age, sex and aged matched for each independent experiment.

### Infections and injections

*Heligmosomoides polygyrus* and *Toxoplasma gondii* infections were done by oral gavage using 200 L3 larvae or 10 cysts of *T. gondii* obtained from either chronically infected C57Bl6 (Vancouver) or CD1 mice (Glasgow, University of Strathclyde), respectively. Use of mice at the University of Strathclyde conformed to guidelines from the Home Office of the UK Government and was covered by the Home office licence PPL60/4568, ‘Treatment & prevention of Toxoplasmosis’. Influenza A virus PR8 strain was administered intranasally to mice anesthetized with isoflurane. Mice were infected with 5 pfu of virus in 25ul. *Nippostrongylus brasiliensis* infection was done by sub-cutaneous injection of 250 L3 larvae. For 2-NBDG uptake, mice were injected intravenously with 100μg of 2-NBDG diluted in sterile PBS 1 hour prior to sacrifice.

### Antibodies and reagents

Staining for flow cytometry was done using the following anti-mouse mAbs: CD3 (17A2) conjugated to PE/Dazzle 594 (BioLegend), CD4 (RM4-5) conjugated to e450, APC APC-Cy7 or Spark YG593 (BioLegend); CD44 (IM7) conjugated to FITC, APC, PE-Cy7 (BioLegend) or BUV737 (BD Biosciences); TCRβ (H57-597) conjugated to PerCP-Cy5.5 (Biolegend); ST2 (DIH9), conjugated to BV421, PE or BV711 (BioLegend); PD-1 (29F.1A12) conjugated to PE, BV711 or BV785 (BioLegend); IL-4 (11B11) conjugated to BV421 (BioLegend); IL-5 (TRFK5) conjugated to PE (BioLegend); IL-13 (eBio13A) conjugated to e660 or PE-Cy7 (ThermoFisher); Foxp3 (FJK-16s) conjugated to e450 or APC (ThermoFisher); GATA3 (TWAJ) conjugated to PE or PerCP-eF710 (ThermoFisher); phospho-S473 Akt (M89-61) conjugated to PE (BD Bioscience); phospho-S235/236 S6 (N7-548) conjugated to Alexa Fluor 647 (BD Bioscience); CXCR5 (L138D7) conjugated to BV650 (BioLegend); Ki67 (16A8) conjugated to BV605 (BioLegend). Separation of dead cells was achieved using e506 or e780 fixable eViability dye (ThermoFisher) or Zombie NIR (BioLegend). 2NBDG and BODIPY FL C16 were purchased lyophilized from ThermoFisher. 2NBDG (2-Deoxy-2-[(7-nitro-2,1,3-benzoxadiazol-4-yl)amino]-D-glucose) was reconstituted in PBS. BODIPY FL C16 (4,4-Difluoro-5,7-Dimethyl-4-Bora-3a,4a-Diaza-*s*-Indacene-3-Hexadecanoic Acid) was reconstituted by shaking for 1 hour in 1% fetal bovine serum (FBS) in PBS. 2-deoxy-d-glucose (2DG) (Sigma-Aldrich) was reconstituted in base DMEM (Aligent).

### In vitro CD4^+^ T cell polarization

CD4^+^ T cells were isolated from naïve splenocytes using the EasySep mouse CD4^+^ T cell enrichment kit (StemCell Technologies) and plated at 3x10^5^ cells per well in a flat bottom 96 well plate. Cells were cultured in complete RPMI (cRPMI) media: RPMI 1640, 10% FBS, 1mM Glutamax, 1mM penicillin-streptomycin, 1mM non-essential amino acids, 1mM sodium pyruvate, and 50μM 2-mercaptoethanol (ThermoFisher). Stimulation was done using 1μg/ml plate-bound αCD3 and 1ug/ml soluble αCD28 (eBioscience) in polarizing conditions; Th1: 10ng/ml IL-12 (ThermoFisher), 10ng/ml IL-2 (ThermoFisher), 1ug/ml αIL-4 (UBC Ablab/BioLegend); or Th2: 40ng/ml IL-4 (ThermoFisher), 10ng/ml IL-2 (ThermoFisher), 1μg/ml αIFNγ (UBC Ablab/BioLegend). Where stated, cells were treated with 0.5mM 2DG. Activated cells were sorted or analysed 4 days after stimulation.

### Lung harvest and digestion

Mice were euthanized by overdose of intraperitoneally injected anaesthetic and lungs were perfused with ice-cold PBS before being removed. Lungs were transferred to 1ml of digest buffer and cut into 1mm pieces. Digest buffer was made by reconstituting 5mg of Liberase TL (Roche) in 2ml of water and using it at a 1/6.5 dilution (∼2U/ml) in HBSS containing Mg^2+^ and Ca^2+^ (ThermoFisher), and 50U/ml of DNAse I from bovine pancreas (Sigma-Aldrich). Cut lung pieces were digested for 35min at 37°C shaking at 185rpm. Digestion was halted by added 5ml of ice-cold cRPMI before crushing digested tissue through a 70μm strainer and washed through with 25ml of cold cRPMI. Cells were spun down at 400xg for 5 minutes and resuspended in LCK buffer (ThermoFisher) for 3 min to lyse red blood cells, before being washed with an additional 25ml of cRPMI. Cells were spun and resuspended appropriately for downstream analysis.

### Ex vivo CD4^+^ isolation and restimulation

Lymph nodes from infected mice were directly crushed into a single-cell suspension through a 70μm strainer in cRPMI. Lung T cells were isolated via tissue digestion as outlined above. Single-cell suspensions were enriched for CD4^+^ cells using either the EasySep mouse CD4^+^ T cell enrichment kit (StemCell Technologies), or MojoSort^TM^ CD4 T cell isolation kit (BioLegend), before being purified by FACS. Sorted cells were plated in 96 well plates as indicated: αCD3, 1μg/ml plate-bound; IL-33, 20ng/ml (BioLegend); PD-L1-Fc, 5μg/ml plate-bound (R&D Systems); Orlistat, 50uM (Tocris). For intracellular cytokine staining, total tissue homogenate/digest was directly re-stimulated with 50 ng/ml phorbol-13-myristate-12-acetate (PMA) (Sigma-Aldrich) and 0.5 μg/ml ionomycin (Sigma-Aldrich) for 5 hours in the presence of 1x monensin (Thermo Fisher). Glucose titrations were done using using glucose-free RPMI-1640 and dialyzed serum (ThermoFisher) plus supplements. cRPMI made with charcoal-stripped serum (ThermoFisher) was used as fatty-acid free media.

### Flow cytometry and cell-sorting

*In vitro* cultures were directly stained with fixable eViability dye (ThermoFisher) and antibodies against surface antigens in FACS buffer for 30 minutes at 4°C. Tissue samples being directly analysed *ex vivo* were Fc-blocked by incubation with 5μg/ml αCD16/32 (BioLegend) for 10 minutes prior to viability and surface staining. Cells were fixed and permeabilized with BD Cytofix for 20 minutes and stained for intracellular cytokines either at 4°C overnight or for at least 1 hour at room temperature. Cells were washed with eBioscience Foxp3 perm/wash buffer (ThermoFisher) and resuspended in FACS buffer for analysis. To maintain reporter expression, transcription factor staining was achieved by fixing cells with BD Cytofix for 1 hour at room temperature after surface staining. Cells were then washed with eBioscience Foxp3 perm/wash buffer and stained overnight at 4°C, or at room temperature for a minimum of 2 hours. Phosflow staining was done using Perm Buffer III (BD Biosciences) according to the manufacturer’s protocol. *Ex vivo* phos-flow analysis was done by resuspending MLN single-cell suspensions directly in pre-warmed 1x fixation buffer. Data for stained cells was acquired using an LSRII (BD Bioscience) in Vancouver and either a BD LSRII or BD Fortessa in Glasgow. Cell sorting was done using a BD Influx or BD Aria II. CD4^+^ T cells were sorted into 2ml of cRPMI using a 100μm nozzle. Analysis was done using FlowJo (TreeStar). Data for the metabolic flow-cytometry panel was obtain and analysed as previously described^25^.

### RNA isolation and quantitative-PCR

RNA was extracted from sorted cells using the RNeasy mini kit (Qiagen) or RNAqueous Micro Kit (Ambion). Concentrations of RNA were determined using a nanodrop 1000. 0.1-1.0μg of RNA was converted to cDNA using the iScript cDNA synthesis kit (Bio Rad) or the High-Capacity RNA-to-cDNA kit (ThermoFisher). qPCR was done using either SsoFast EvaGreen supermix and a BioRad CFX96 Real Time System (Vancouver), or PowerUp SYBR Green master mix (ThermoFisher) using a QuantStudio 6 Flex Real-time PCR system (Applied Biosystems) (Glasgow). Target gene expression was normalized to the expression of the ribosomal protein S29 (*RPS29*, Fwd 5’-ACGGTCTGATCCGCAAATAC-3’, Rev 5’-CATGATCGGTTCCAC TTGGT-3’) using the △Ct method. *HKII, ENO1, PKM2* and *LDHA* primer sequences were obtained from Shi *et al*, 2011^6^; *PPARG* sequences from Nobs *et al,* 2017 ^34^; *CD36* sequences from Matsusue *et al,* 2014^57^; *CPT1A* sequences from Byersdorfer *et al*, 2013^58^; *FABP4* and *FABP5* from Pan *et al*, 2017^59^; *FASN, ACLY, ACACA* from Young *et al*, 2017^60^. Primers were order from ThermoFisher.

### Seahorse metabolic flux assay

2x10^5^ sorted *in vitro* or 4x10^5^ sorted *ex vivo* activated or naive cells were plated in glucose-free minimal DMEM (Aligent Technologies), incubated for 1 hour at 37°C, and analysed using a Seahorse XFe96 Bioanalyzer. ECAR and OCR were measured following injection of 10mM glucose, 2μM oligomycin, and 50mM 2DG (Sigma-Aldrich). Results are shown normalized to cell number. Calculations were performed using the greatest values of individual wells after each injection.

### Metabolite uptake and mitochondrial staining

To determine 2NBDG and BODIPY FL C16 uptake single-cell suspensions from harvested tissues pooled and CD4^+^ T cells were purified using the MojoSort CD4^+^ isolation kit. A 96 well round-bottom plate was seeded with 2.5x10^5^ purified CD4^+^ cells and rested in cRPMI for 30 minutes at 37°C. Cells were spun down and resuspended in 50μM 2NBDG or 5uM BODIPY FL C16 diluted in PBS and incubated at 37°C for 15 minutes before being quenched with ice-cold FACS buffer, washed twice, and stained for surface markers. To stain for mitochondria, single-cell suspensions were incubated for 20 minutes with 50nM of MitoSpy Orange CMTMRos (BioLegend) and 50nM of MitoTracker DeepRed FM (ThermoFisher) in RPMI containing no additives. Cells were washed twice with PBS before additional staining.

## Statistical analysis

Statistics were assessed using GraphPad Prism version 8. For direct comparison between 2 groups, an unpaired students T test was used. For comparisons between 3 or more groups, a one-way analysis of variance (ANOVA) was performed with Tukey’s multiple comparison correction. In cases where groups had unequal standard deviations, unpaired T-tests were run with Welch’s correction or Brown-Forsythe and Welch ANOVA was used with the Dunnett T3 correction for multiple comparisons. Data are presented as the mean with standard deviation. \**P* ≤ 0.05, \*\**P* ≤ 0.01, \*\*\**P* ≤ 0.001, \*\*\*\**P* ≤ 0.0001, ns = not significant.

## Acknowledgements

We thank David Dow, Joanne Battersby and staff in the Wolfson Research Unit for animal husbandry; Nicola Britton and Claire Ciancia for *H. polygyrus* larvae and life cycle maintenance; the Flow Core at the University of Glasgow and especially Diane Vaughan for flow cytometry support; as well as Andy Johnson and Justin Wong of the Flow Core at the University of British Columbia. This work was supported by the MRC (grant MR/S009779/1), the CIHR (MOP-126061), and funds from the University of Glasgow and the University of British Columbia awarded to GPW. RMM also received Wellcome Trust support through an Investigator Grant (Ref 219530), and the Wellcome Trust core-funded Wellcome Centre for Integrative Parasitology (Ref: 104111). GAH was supported by a Natural Sciences and Engineering Research Council postgraduate doctoral scholarship (grant 653357) and UBC four-year fellowship, and was awarded a Leiden University Funds grant (www.luf.nl, W213032-2-38).

## Author Contributions

G.A.H and G.P.W. developed the project and wrote the manuscript. G.A.H. carried out experiments and analysis. R. M. M. provided reagents, helminth larvae, and project insight. B.E. provided reagents and project insight. G.P.W acquired funding and supervised the study.

## Declaration

The authors have no conflicts of interest to declare.

---

**Figure S1:**
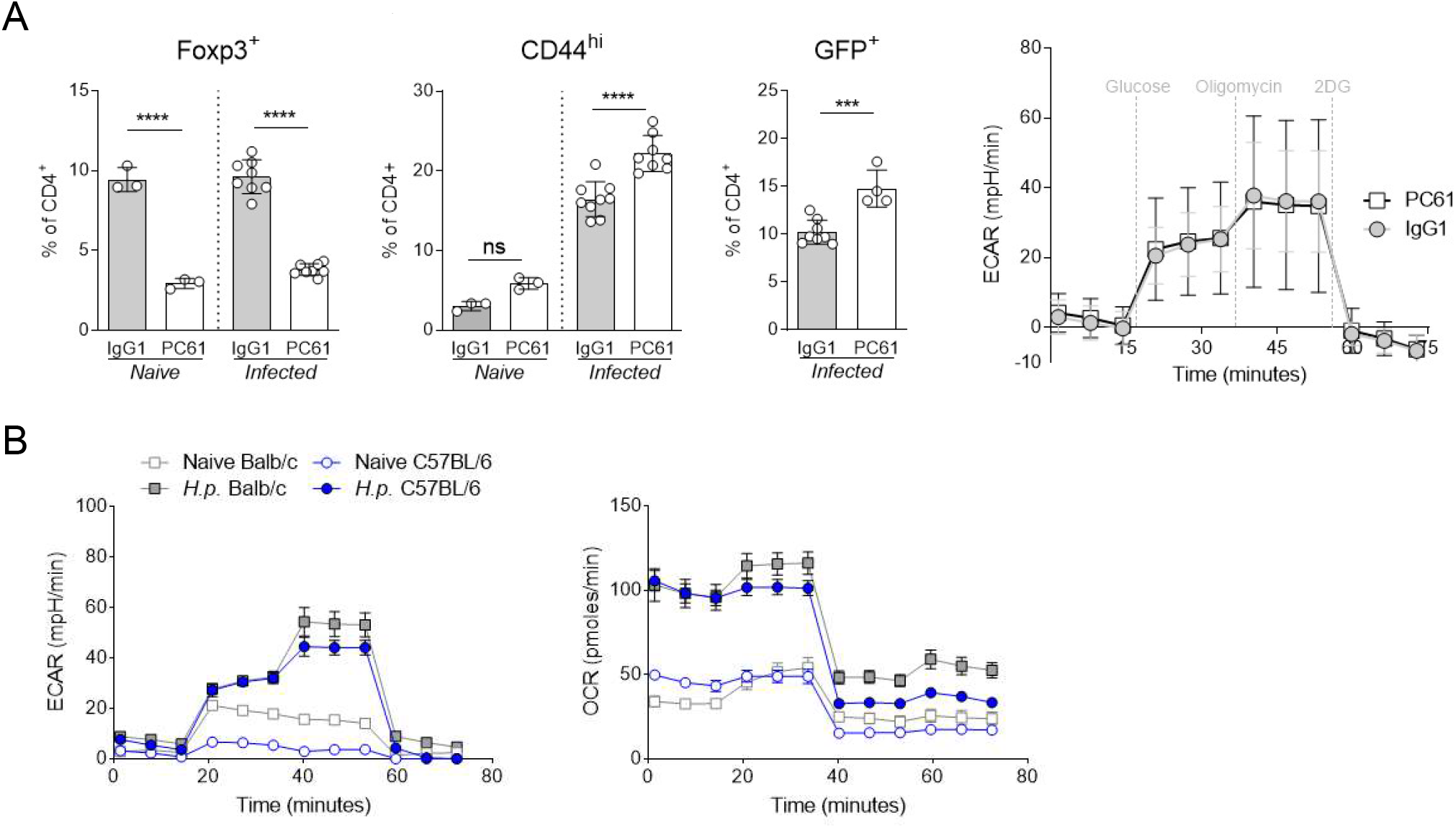
Th2 metabolism is independent of Tregs for susceptibility to infection. (A) Frequency of Foxp3^+^ Tregs, activated CD4^+^CD44^Hi^ and il4-expressing T cells at day 14 of Hp infection, and glycolysis stress test of sorted *il4^+^* cells from the MLN, following a single dose of 1μg anti-CD25 depleting antibody (PC61) one day prior to the start of infection; representative of 2 experiments with individual mice shown as separate data points, or pooled for technical replicates for Seahorse analysis. (B) Seahorse traces from glycolysis stress test of Th2 cells sorted from the MLN of either Balb/c or C57BL/6 background reporter mice. Students’ two-tailed t-test used for statistical analysis: ****p<0.0001, ***p<0.001.

**Figure S2:**
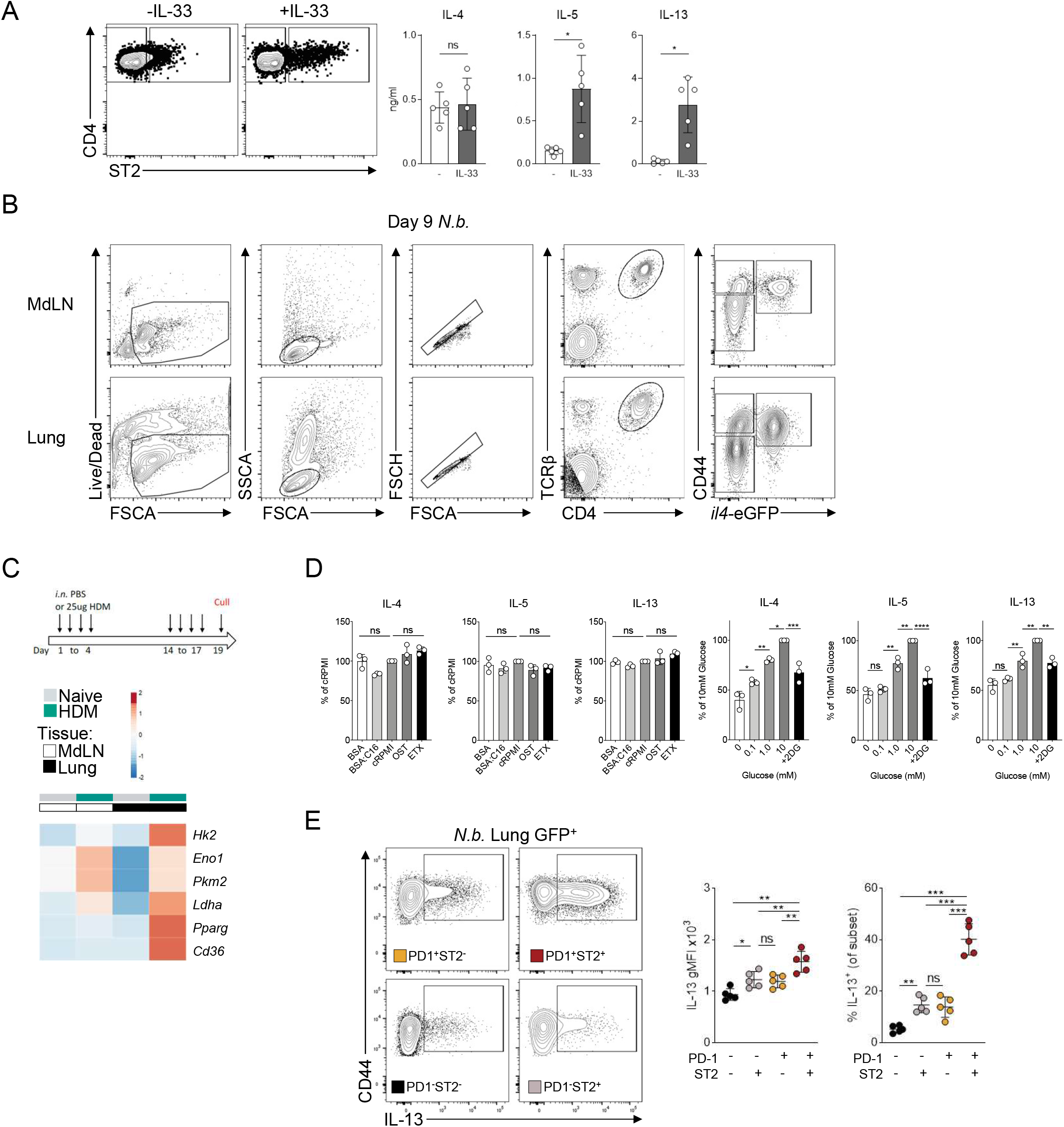
FA metabolism in Th2 cells does not directly impact cytokine production. (A) ST2 expression of sorted MLN CD44hi cells cultured overnight after CD3 stimulation in the presence of absence of IL-33, and corresponding cytokine production measured from the supernatants by cytometric bead array. (B) Gating strategy for identifying Th2 cells from Nb infected mice. (C) Relative expression of metabolic targets from sorted Th2 cells from unchallenged or HDM sensitized and challenged mice, showing mean from 3 mice. (D) Cytokines detection in GFP+ Th2 cells after direct ex vivo stimulation of total lung homogenate in the presence or absence of FA (using charcoal-stripped serum +/- BSA:palmitate), CPT1A inhibitor etomoxir, lipase inhibitor orlistat, increasing glucose concentrations, or Hk2 inhibitor 2-DG; data points are individual mice, representative of 2 experiments. (E) Intracellular IL-13 expression after PMA/ionomycin stimulation in PD-1 and ST2 expressing populations of lung Th2 cells, corresponding to Figure 2F. Statistics analysed by student’s two-tailed t-test or one-way ANOVA: ****p<0.0001, ***p<0.001, **p<0.01, *p<0.05, ns = not significant.

**Figure S3:**
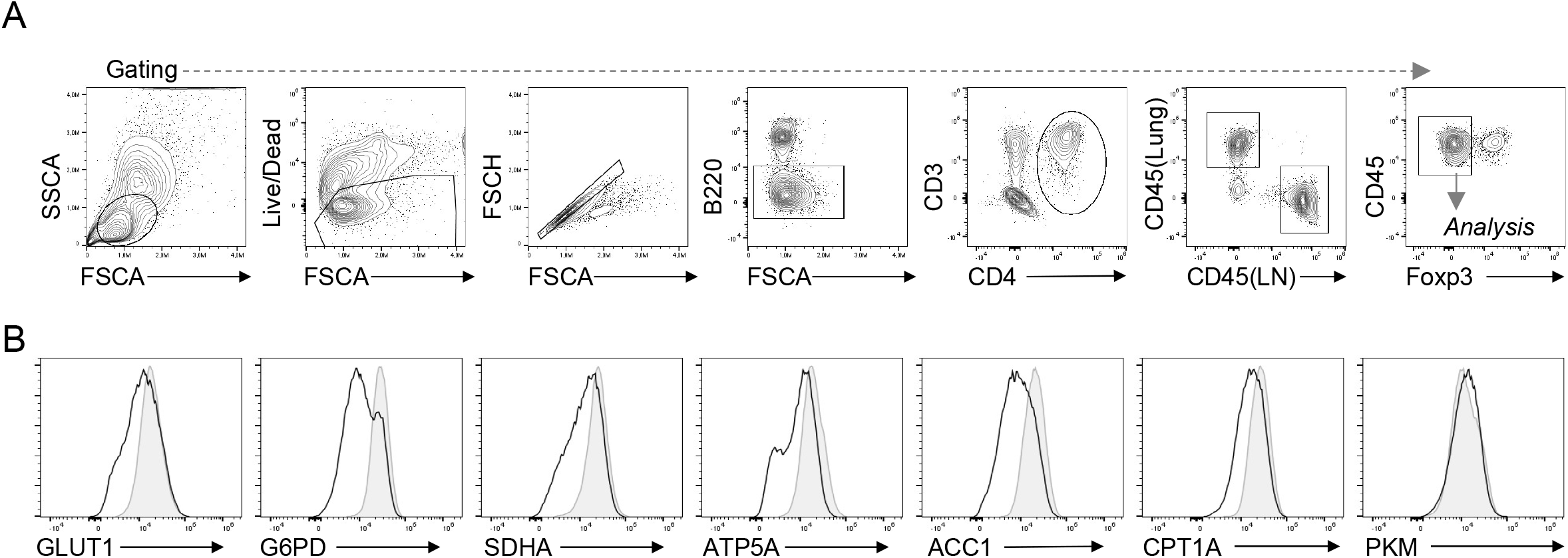
Gating for metabolic panel. (A) Representative gating for barcoded and pooled MdLN and Lung samples from Nb infected mice. (B) Representative histograms for metabolic targets for MdLN (grey) and Lung (black).

